# Lognormal Neural Point Process Models for Interpretable Heartbeat Dynamics

**DOI:** 10.64898/2026.08.12.744524

**Authors:** Bhuvana Ravi Kumar, Bharath Ramsundar, Sandya Subramanian

## Abstract

Neural temporal point processes (NTPPs) are powerful tools for modeling sequences of timestamped events with statistical temporal structure. Density-based NTPPs, in particular, are an interesting opportunity to merge the universal function approximation capability of neural networks with a defined statistical model in a way that has many potential applications. We demonstrate one such application to heartbeat dynamics, a physiologic point process. We specifically apply a lognormal mixture NTPP to compute instantaneous estimates of the mean and standard deviation of beat-to-beat intervals. We compare our results to the state of art (Barbieri et al.) point process model for heartbeat dynamics, which uses a more physiologically rigorous inverse Gaussian model. We find that the NTPP model maintains reasonable accuracy while improving upon robustness to noise.

## 1 Introduction

NTPPs (4) offer a flexible framework that blends traditional statistics and AI to to model event intervals in temporal sequences. However, they remain underutilized in many domains such as physiology despite the presence of known generative structures. Heartbeats, for instance, arise from ions flowing across cell membranes in the heart and causing electrical currents. Heartbeat intervals, termed RR intervals, are known to have inverse Gaussian structure (2). Barbieri et al. (2) proposed a parametric model that while grounded in this physiology, can be highly sensitive to noise and artifact and computationally expensive. In previous work, we applied point-process goodness of fit frameworks to rigorously validate NTPP fit to heartbeat dynamics (1). In this work, we show that a lognormal mixture NTPP with GRU-based history encoding preserves statistical structure and physiologic interpretability while improving computational performance and robustness. Importantly, the NTPP model retains the principle of history-dependent RR interval prediction, central to Barbieri’s approach, while enhancing computational tractability through efficient parallelizable inference.

## 2 Methods

We used 4 hours of continuous ECG-derived RR interval data from each of 10 healthy volunteers. The NTPP model is a lognormal mixture model with a GRU-based encoder (1). We employed a leave-one-subject-out training strategy: for each test subject, the model was trained on 8 other subjects, validated on 1 held-out subject, and evaluated on the remaining subject. In this work, we report results on the final test subject only. For comparison, we computed Barbieri model estimates of instantaneous mean and standard deviation of the RR interval on the same test dataset. These served as the physiological benchmark. Barbieri’s history-dependent inverse Gaussian model estimates the instantaneous RR interval distribution conditioned on past heartbeats assuming a linear autoregressive structure (2). The NTPP model outputs a lognormal mixture density over RR intervals, in which each component *k* has a log-mean *μ*_*k*_, log-std *σ*_*k*_, and weight *w*_*k*_. From these, we compute:

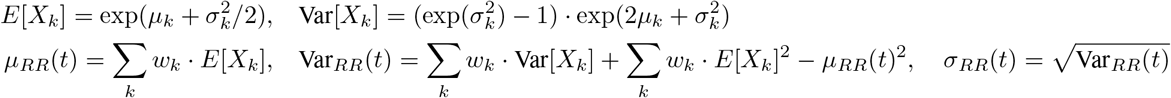

## 3 Results

Figure 1 shows the NTPP model’s statistical goodness-of-fit over the training process and time-rescaling validation (1).

**Figure 1:**
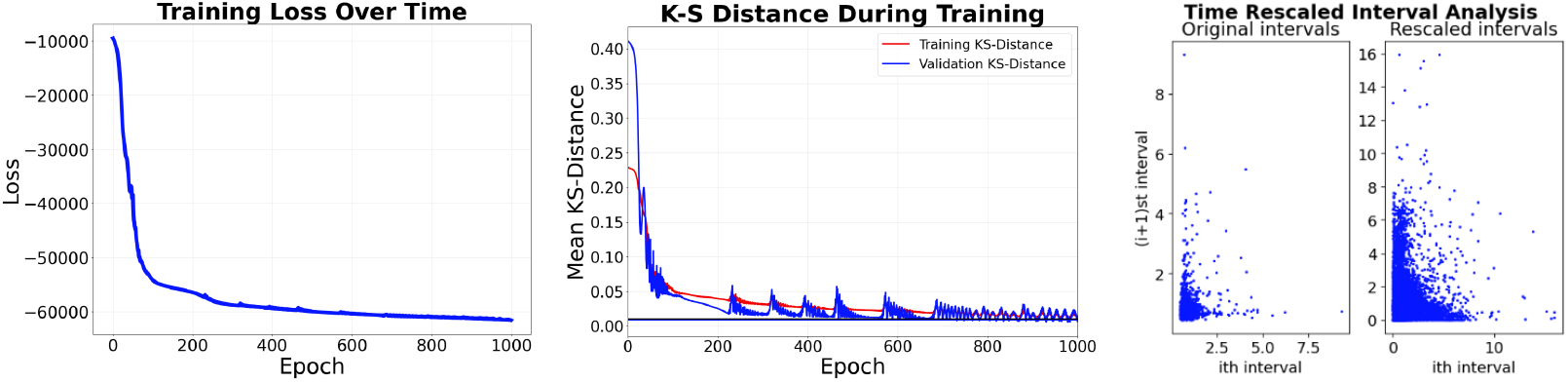
Model training and statistical validation: Left – loss convergence; Middle – decreasing KS-distance over epochs; Right – time-rescaled RR intervals showing near-uniformity, confirming model fit.

Figure 2 compares the instantaneous estimates of the mean and standard deviation of the RR interval between the NTPP model and the Barbieri model, as well as the actual RR intervals.

**Figure 2:**
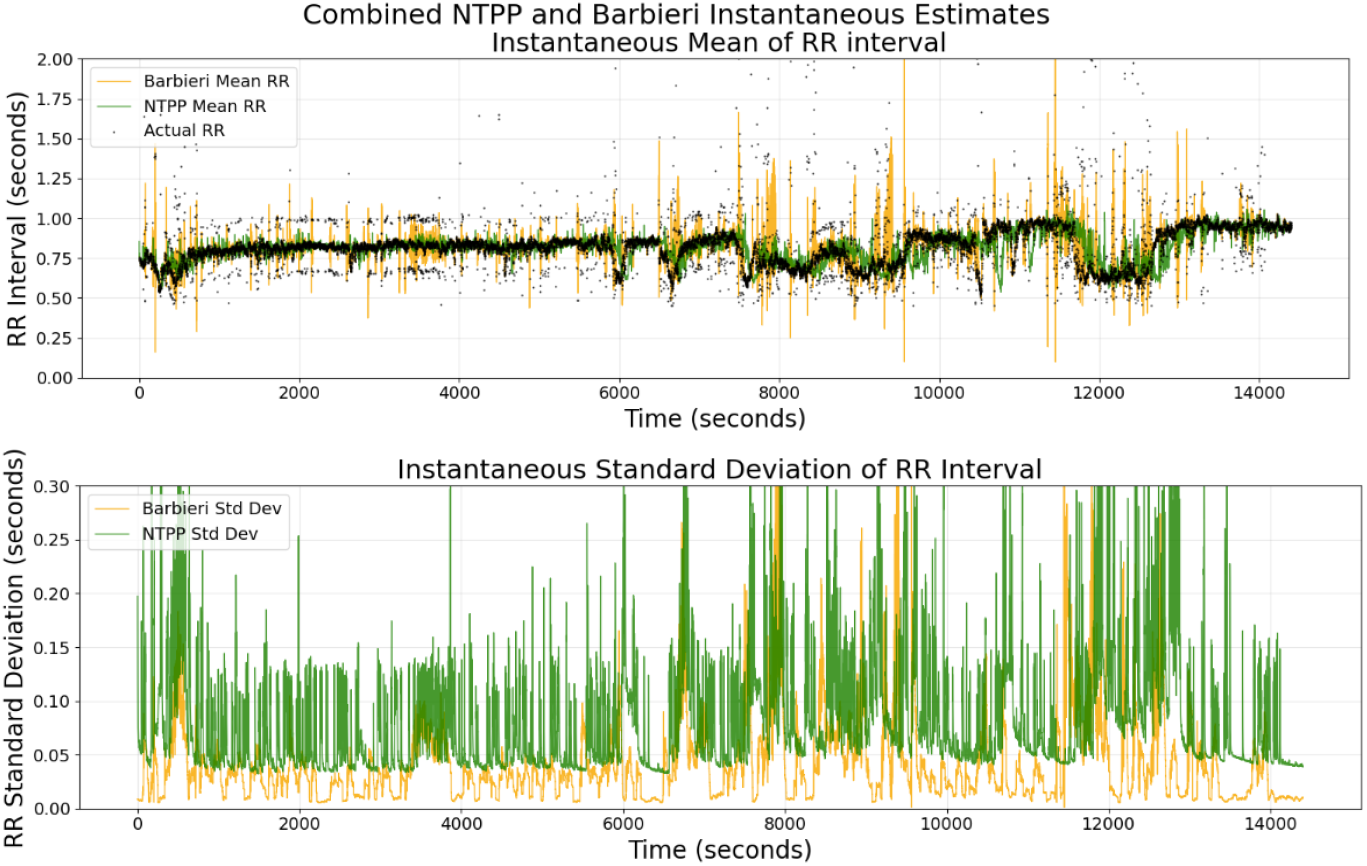
Instantaneous RR mean and standard deviation. NTPP (green) better tracks ground truth (black) than Barbieri’s model (orange), though it predicts higher variability.

## 4 Discussion

The NTPP model is a reasonable fit to the data because the KS distance decreases and the correlation between the rescaled intervals is more scattered (Fig. 1). The NTPP model more closely follows the actual RR intervals than the Barbieri model, largely due to increased robustness to noise and outliers in the identified R peaks (Fig. 2). However, the NTPP model’s standard deviation estimates are consistently higher, due to heavier tails in the lognormal mixture compared to the inverse Gaussian distribution. This can be improved by replacing the lognormal mixture with an inverse Gaussian mixture, which is also more physiologically rigorous. This case study illustrates the potential of density-based NTPPs to model physiologic phenomena like heartbeat dynamics with statistical rigor and computational tractability. Compared to classical point process models, NTPPs offer greater modeling flexibility and improved computational efficiency without sacrificing interpretability. This makes them well suited to modeling several other natural phenomena that are point processes, such as volcanic eruptions and neural spikes.

